## Supplementary File for "The autophagy protein, ATG14 safeguards against unscheduled pyroptosis activation to enable embryo transport during early pregnancy"

**This PDF file includes:**

**1. Supplementary Figure 1:** *Atg14* cKO mice show normal gross morphology of uterus and ovary.

**2. Supplementary Figure 2:** *Atg14* cKO mice altered autophagy markers expression.

**3. Supplementary Figure 3:** *Atg14* cKO mice show a reduced number of FOXJ1-expressing ciliary epithelial cells.

4. **Supplementary Figure 4**: Average number of embryos are unaltered in *Atg14* cKO or Polyphyllin-treated D4 pregnant mice

**3. Supplementary Table 1:** List of primers and TaqMan probes

**4. Supplementary Table 2:** List of antibodies

**Supplementary Figure Legends:**

**Figure S1: *Atg14* cKO mice show normal gross morphology of uterus and ovary.** Histological analysis using H&E staining of 8-week-old virgin *Atg14* control and cKO mice uteri and ovary. LE: luminal epithelium, GE: glandular epithelium, ST: stroma, AF: antral follicle, CL: corpus luteum.

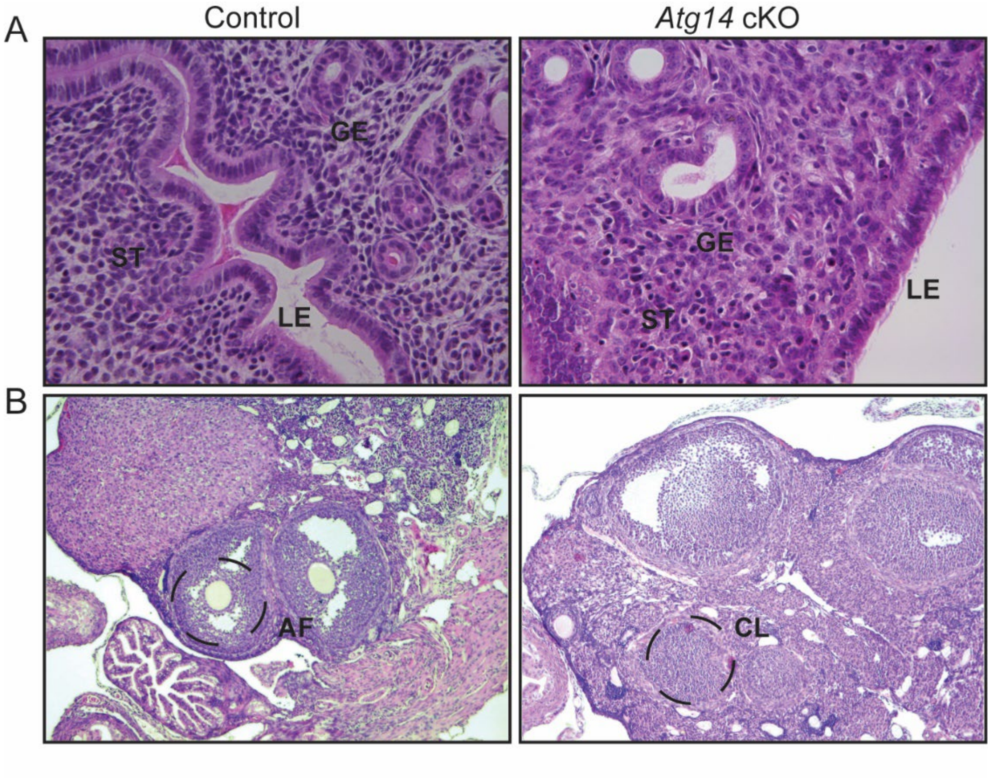

**Figure S2:** ***Atg14*** **cKO mice altered autophagy markers expression.** (**A**) Immunofluorescence analysis to show ATG14 expression in ampulla and isthmus regions of oviducts. LE: Luminal epithelium, M: smooth muscle. Scale bar: 100 μm; 20 μm. The upper panel shows the 10X objective images and the lower panel shows the 20X objective images. (**B**) Western blotting shows LC3B and p62 expression in control and cKO oviducts tissues. β-actin was used as a loading control.

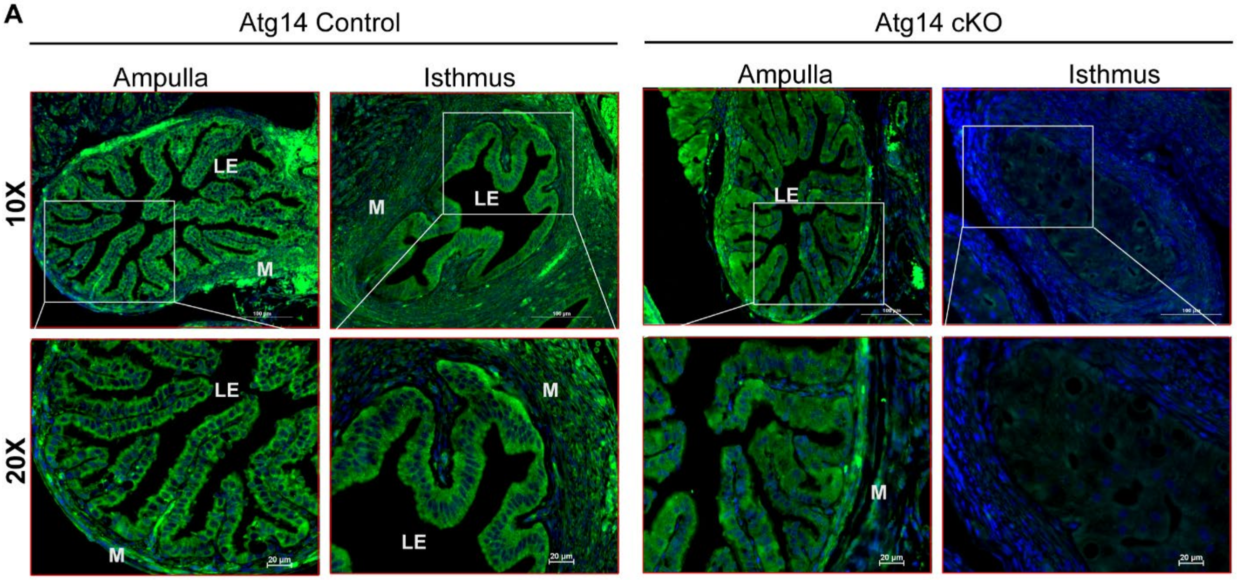


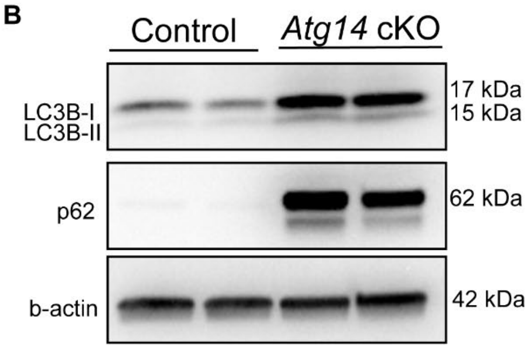


**Fig. S2**

**Figure S3**: ***Atg14* cKO mice show a reduced number of FOXJ1-expressing ciliary epithelial cells.** Immunohistochemistry to show FoxJ1-positive staining in *Atg14* control and cKO oviducts**.** The upper panel shows the ampullary section and the lower panel shows the isthmus section. Images were taken at 20X. Scale bar: 10µm.

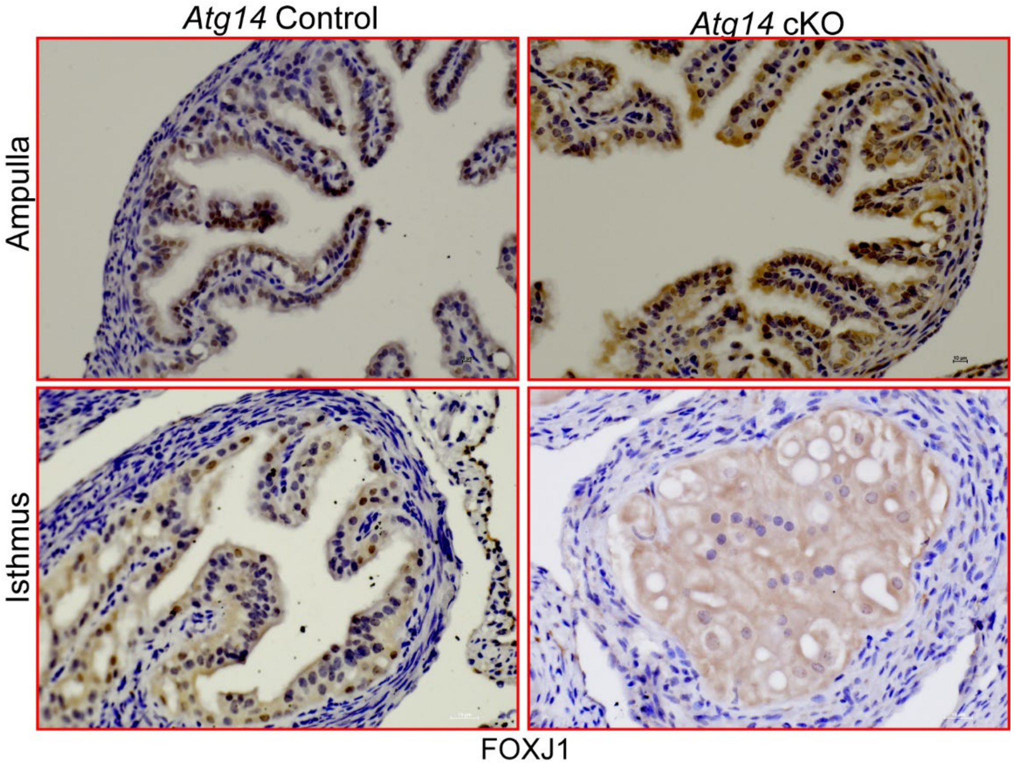


**Fig. S3**

**Figure S4: Average number of embryos are unaltered in *Atg14* cKO or Polyphyllin-treated D4 pregnant mice. (A)** Average number of embryos retrieved from *Atg14* cKO (n=5) oviducts or control uteri (n=5). **(B)** Average number of embryos retrieved from vehicle (n=3) uteri and polyphyllin-treated (n=3) D4 pregnant mice oviducts & uteri.

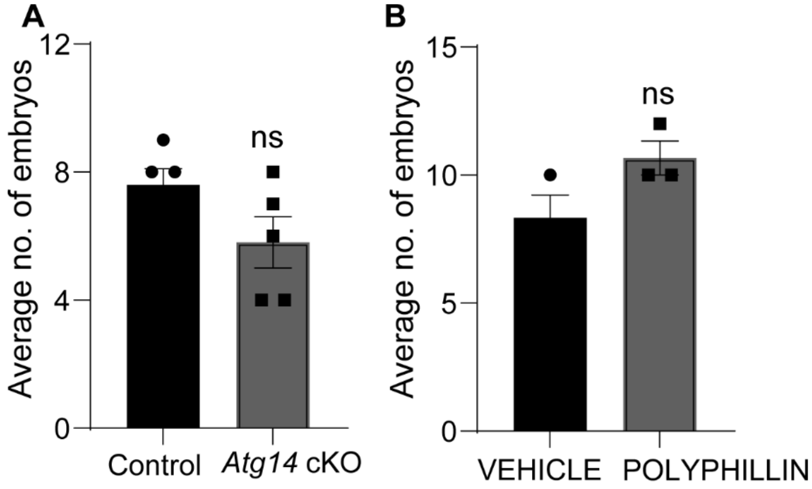


**Fig. S4**

**Supplementary Table 1. List of primers and TaqMan probes**

| Gene name | Species | Application, Chemistry | Company | Sequence/Cat. No. |
|---|---|---|---|---|
| Atg14 f/f | Mouse | Genotyping | IDT | P1: TTGACCGTCACAGGGTGTGAGTGACTT<br>P2: AAGCAGAGTTAGGCTTCCCTGGTAGAA<br>P3: CCCATCTCCATTCCTGGATTACTGGAC<br>P4: CTAAAGCGCATGCTCCAGACTGCCTTG |
| $PR^{Cre}$ | Mouse | Genotyping | IDT | P1: ATGTTTAGCTGGCCCAAA TG<br>P2: TAT ACC GAT CTC CCT GGA CG<br>P3: CCC AAA GAG ACA CCA GGA AG |
| $Foxj1^{Cre}$ | Mouse | Genotyping | IDT | P1: ATTTGGGCCAGCTAAACATGC<br>P2: GCAAAACAGGTAGTTATTCGG |
| *Esr1* | Mouse | qPCR, TaqMan | ABI | Mm00433149_m1 |
| *Pgr* | Mouse | qPCR, TaqMan | ABI | Mm00435628_m1 |
| *Atg14* | Mouse | qPCR, TaqMan | ABI | Mm00553733_m1 |
| *Tnf-alpha* | Mouse | qPCR, TaqMan | ABI | Mm00443258_m1 |
| *Cxcr3* | Mouse | qPCR, TaqMan | ABI | Mm99999054_s1 |
| *Lif* | Mouse | qPCR, TaqMan | ABI | Mm00434761_m1 |
| *Mcm2* | Mouse | qPCR, TaqMan | ABI | Mm00484804_m1 |
| *Ccnd1* | Mouse | qPCR, TaqMan | ABI | Mm00432359_m1 |
| *Fgf18* | Mouse | qPCR, TaqMan | ABI | Mm00433286_m1 |
| *ATG14* | Human | qPCR, TaqMan | ABI | Hs00208732_m1 |
| *18S* | Human<br>Mouse | qPCR, TaqMan | ABI | 4318839 |

*All primer sequences are written 5' to 3'

ABI-applied biosystems

IDT-integrated DNA technologies.

**Supplementary Table 2. List of antibodies**

| Antibody | Company, Catalogue number | Application |
|---|---|---|
| ATG14 | Proteintech, 24412-1-AP | Immunofluorescence |
| GSDMD | Thermo Fisher Scientific, Cat# PA5-115330 | Immunoblotting |
| GSDMD | Abcam, ab209845 | Immunofluorescence |
| CASPASE 1 | Abcam, ab138483 | Immunoblotting |
| Ki-67 | Abcam, ab15580 | Immunofluorescence |
| FOXJ1 | Sigma, HPA 005714 | Immunohistochemistry |
| PAX8 | CST, #59019s | Immunohistochemistry |
| MUC1 | Abcam, ab15481 | Immunofluorescence |
| TOM20 | ab186735 | Immunofluorescence |
| CYTOCHROME C (6H2.B4) | Thermo Fisher Scientific, 33-8200 | Immunofluorescence |
| Alpha-smooth muscle actin (KRT8) | Developmental Studies Hybridoma Bank, TROMA-I | Immunofluorescence |
| Normal Rabbit IgG | CST, #2729 | Immunofluorescence |
| Goat anti-Rat IgG (H+L) Cross-Adsorbed Secondary Antibody, Alexa Fluor™ 488 | Thermofisher Scientific, A11006 | Immunofluorescence |
| Goat anti-Rabbit IgG (H+L) Highly Cross-Adsorbed Secondary Antibody, Alexa Fluor™ 488 | Thermofisher Scientific, A11034 | Immunofluorescence |
| Goat anti-Rabbit IgG (H+L) Highly Cross-Adsorbed Secondary Antibody, Alexa Fluor™ 594 | Thermofisher Scientific, A11037 | Immunofluorescence |
| GAPDH | CST, #2118S | Immunoblotting |
| Anti-rabbit IgG, HRP-linked | CST, #7074 | Immunoblotting |
